# Annexin-A1 tripeptide attenuates surgery-induced neuroinflammation and memory deficits through regulation of the NLRP3 inflammasome

**DOI:** 10.1101/2020.05.12.090654

**Authors:** Zhiquan Zhang, Qing Ma, Ravikanth Velagapudi, William E. Barclay, Ramona M. Rodriguiz, William C. Wetsel, Mari L. Shinohara, Niccolò Terrando

**Author notes:** correspondence: Zhiquan Zhang, PhD; Niccolò Terrando, PhD.

## Abstract

Neuroinflammation is a growing hallmark of perioperative neurocognitive disorders (PNDs), including delirium and longer-lasting cognitive deficits. We have developed a clinically-relevant orthopedic mouse model to study the impact of a common surgical procedure on the vulnerable brain. The mechanism underlying PNDs remain unknown. Here we evaluated the impact of surgical trauma on the NLRP3 inflammasome signaling, including the expression of apoptosis-associated speck-like protein containing a CARD (ASC), caspase-1, and IL-1β in the hippocampus of C57BL6/J male mice, adult (3-months) and aged (>18-months). Surgery triggered ASC specks formation in CA1 hippocampal microglia, but without inducing significant morphological changes in NLRP3 and ASC knockout mice. Since no therapies are currently available to treat PNDs, we assessed the neuroprotective effects of a biomimetic peptide derived from the endogenous inflammation-ending molecule, Annexin-A1 (ANXA1). We tested the hypothesis that this peptide (ANXA1sp) inhibits NLRP3 inflammasome activation, thus preventing microglial activation and hippocampal-dependent memory deficits. Together these results uncover a previously underrecognized role of the NLRP3 inflammasome in triggering postoperative neuroinflammation and offer a new target for advancing treatment of PNDs through resolution of inflammation.

## Introduction

Perioperative neurocognitive disorders (PNDs), which include acute delirium and lingering cognitive impairment following surgery and hospitalization, are common especially within the rapidly growing senior population (Evered et al., 2017). These neurologic complications are significant problems that can profoundly alter recovery from routine procedures such as orthopedic surgery (Subramaniyan and Terrando, 2019). Aside from the implications of a worsened perioperative course that include prolonged hospital length-of-stay, higher rates of nursing home placement, and soaring healthcare costs, PNDs lead to a persistent functional decline, including dementia, and they impact also mortality rates (Monk et al., 2008; Fong et al., 2015). Although some strategies, such as unit-based targeted multifactorial intervention and proactive geriatric consultation have been implemented to prevent postoperative delirium (Marcantonio, 2017). Importantly, there is no Food and Drug Administration approved drug available to treat PNDs

Aberrant inflammation is generally considered to be a driver of PNDs (Safavynia and Goldstein, 2018). We have developed a clinically-relevant mouse model of PNDs by using a tibial fracture method that mimics common aspects of orthopedic trauma and repair, and we have studied its effects on the central nervous system (CNS) (Cibelli et al., 2010; Terrando et al., 2010; Vacas et al., 2014). Using this model, we have demonstrated a role for systemic cytokines that promote neuroinflammation and glial activation in the hippocampus (Cibelli et al., 2010; Terrando et al., 2010). Anti-inflammatory agents can ameliorate this cytokine storm and, thereby prevent neuroinflammation and the presentation of PND-like behaviors in rodents. However, the possibility of side effects that involve overt immunosuppression, raises concerns for clinical translation (Simone and Tan, 2011). Resolution pharmacology has provided a shift in the approach to treat inflammatory conditions by focusing on stimulating pro-resolving pathways *versus* simply blocking pro-inflammatory molecules (Perretti et al., 2015). To date several endogenous pro-resolving pathways have been identified and these include bioactive lipids, autacoids, and peptides (e.g., annexin-A1 or ANXA1) (Serhan and Levy, 2018). These bioactive mediators have been implicated in arthritis, diabetes, stroke, dementia, and other inflammatory conditions, yet little is known about their mechanisms of action and targets in PNDs.

ANXA1 is a glucocorticoid-regulated protein with well-established effects on the immune system, that involve regulation of leukocyte trafficking, endothelial integrity, and microglial activation (Gavins and Hickey, 2012). Several peptides (Ac2-26, Ac2-12, and Ac2-6) have been derived from the full-length N-terminal domain of ANXA1 and they have been tested in several inflammatory conditions (Sheikh and Solito, 2018). We have developed a small peptide (ANXA1sp) with potent anti-inflammatory effects on NF-κB signaling and organ-preserving functions (Zhang et al., 2010; Zhang et al., 2017; Ma et al., 2019). Based on preclinical and clinical results on the involvement of interleukin (IL)-1β signaling in PNDs (Cibelli et al., 2010; Cape et al., 2014), we have hypothesized that NLRP3 inflammasome activation orchestrates postoperative neuroinflammation. Here, we tested the ability of ANXA1sp to modulate NLRP3 inflammasome activation and show that orthopedic surgery induces ASC speck formation in the hippocampus, suggesting a role for the NLRP3 inflammasome complex in microglia that may control the development of PND-related behavior.

## Materials and Methods

### Animals

The experimental protocol was approved by the Duke University Animal Care and Use Committee. All procedures were conducted under protocols approved by the Institutional Animal Care and Use Committee at Duke University in accordance with the guidelines of the National Institutes of Health for animal care (Guide for the Care and Use of Laboratory Animals, Health and Human Services, National Institutes of Health Publication No. 86-23, revised 1996). Studies were conducted on C57BL6/J male adult mice (3 months; weight 20-26 g) and aged mice (18-22 months; 26-55 g) (Jackson Laboratories, Bar Harbor, Maine). *Nlrp3*^−/−^, *Asc*^−/−^ and ASC-citrine mice (3-4 months; weight 20-26 g) were bred in house. Mice were housed (3-5 animals/cage) in a 12:12-hour light:dark cycle in a humidity- and temperature-controlled room with free access to food and water. The animals were acclimated for at least 1 week before starting any experiment. Note, in the aged cohort, 14 mice were excluded due to complications after surgery (tumor, stroke, death), while no mice from any other cohort were excluded.

### Drug Treatments

Peptide preparation and treatment were performed as previously described (Zhang et al., 2017). Briefly, ANXA1 biomimetic tripeptide (ANXA1sp, Ac-QAW, or QW-3 (Zhang et al., 2010); Ac = acetyl, MW = 445.47 Da), was synthesized and purified (> 98% purity) by GenScript (Piscataway, NJ, USA). The tripeptide was suspended in 100% DMSO and diluted in saline to a final dose of 1 mg/kg ANXA1sp in 0.5% DMSO. Mice were injected intraperitoneally with 1 mg/kg ANXA1sp in 0.5% DMSO and saline 30 minutes before orthopedic surgery.

### Orthopedic Surgery and Sample Preparation

We performed an open tibial fracture as previously described (Velagapudi et al., 2019). Briefly, mice were injected with 0.1 mg/kg buprenorphine (s.c.) and they underwent orthopedic surgery with 2.1% isoflurane anesthesia. Mice not exposed to surgical manipulations served as naïve controls. Mice were euthanized under isoflurane anesthesia and were perfused with ice-cold PBS to flush blood from the brain vasculature. Hippocampi were rapidly (< 2 minutes) dissected on ice, frozen in liquid nitrogen for protein assays, and stored at −80°C until assay.

Mice destined for immunostaining were perfused with an ice-cold solution of 4% paraformaldehyde (PFA) PBS solution. Whole brains were incubated at 4°C in the 4% PFA solution overnight. The next day brains were rinsed in ice-cold PBS and incubated in 20% and 30% sucrose at 4°C for 24 hours, respectively. The brains were placed into OCT solution and stored at −80°C for further use.

### Western blot

Mouse hippocampi were lysed in radioimmunoprecipitation assay (RIPA) buffer (R0278; Sigma) containing Halt Protease Inhibitors Cocktail (78430; Thermo Scientific). Proteins were separated on 4%-15% or 4%-20% Criterion TGX precast protein gels (64134751; Bio-Rad), and transferred to an Immun-Blot PVDF membrane (1620177; Bio-Rad). After blocking with 5% nonfat dry milk (170-6404; Bio-Rad), the blot was probed with the following antibodies: NLRP3 (LS-C374964; LSBio), ASC (67824S; Cell Signaling), Caspase1 (Cleaved Asp210) (LS-C380449; LSBio), IL-1β (LS-C104813; LSBio), and Annexin-A1 (ANXA1) (3299S; Cell Signaling) or (71-3400; Invitrogen). The proteins were visualized using SuperSignal West Dura Extended Duration Substrate (34076; Thermo Scientific) on a ChemiDoc MP Imaging System (Bio-Rad). Protein loading was assessed using anti-GAPDH (2118S; Cell Signaling). Bands were quantified with Fiji software, and analyzed with Prism 8 (GraphPad Software). Hippocampal levels of ATP were determined using a commercially available assay kit (ab83355, Colorimetric/Fluorometric) following the manufacturer’s instructions.

### Immunofluorescence

Immunostaining and analyses of microglial morphology were performed on the PFA-perfused slices using ionizing calcium-binding adaptor molecule 1 (Iba1) rabbit antibody (019-19741; Wako). Briefly, brain slices (30 μm thick) were incubated in 10% normal donkey or goat serum in PBST [1 × PBS (11189, Gibco) + 0.5% Tween 20 (P1379, Sigma)] containing 0.3% Triton × 100 (T8787, Sigma) for antigen retrieval and to block any background for 2 hours at RT. Slices were incubated with the primary antibody (1:500) at 4°C overnight. After 3 washes with PBST, slices were incubated with the secondary antibody conjugated with Alexa Fluor 488 or Fluor 647 (1:500, Invitrogen) for 2 hours at RT. After 5 washes with PBST, the sections were mounted, dehydrated, and cover-slipped. Images were acquired using Zeiss 880 inverted confocal Airy scan.

### Image Analyses

Imaged tile scans were stitched together and presented as high-resolution images. Microglial morphology was quantified as described in (Schwarz et al., 2012), based on 4 morphologic subtypes: round/amoeboid microglia (*Round*), stout microglia (*Stout*), microglia with thick long processes (*Thick*), and microglia with thin ramified processes (*Thin*). The number of Iba1-positive cells was determined in three representative areas of the hippocampal CA1 region per animal. To determine further the state of microglial activation, we manually traced their branch lengths using Imaris Filament Tracer to compare branch lengths from soma and the complexity of individual microglia in the CA1 region before and after orthopedic surgery.

### Behavioral assays: “What-where-when” memory and Memory Load

Hippocampal-dependent memory was assessed with two mnemonic tasks (Huffman et al., 2019; Miller-Rhodes et al., 2019). The first task assessed memory for objects (“what”), their locations (“where”), and the order in which they were presented (“when”). A second task examined memory load with seven unique objects. For the What-Where-When (W-W-W) task, training/testing were conducted in a 65 × 52 × 25 cm arena with complex visual cues on the walls of the room. Briefly, mice were acclimated to the arena for 5 min and then were subjected to two 5-min training and a single test trial, each separate by 55-60 min. In first training trial mice were presented with 4 identical objects (Set A). Three of the objects were spaced equally along the long axis of one wall with the fourth object centered along the opposite wall in a triangle formation. In the second training trial (Set B), a new set of 4 identical objects was presented. Objects 1 and 2 of Set B overlapped in location with objects 1 and 3 of Set A, while the two remaining objects of Set B were positioned in corners 3 and 4. At testing two objects each from Sets A and B were used, for a total of 4 objects. One of the Set A objects was placed in corner 1 (stationary object A). The second Set A object was placed in corner 3 (displaced Set B object). Set B objects remained in corners 2 and 4 (“recent” objects). Behavior was filmed for all trials and scored subsequently with Noldus Ethovision 11 (Noldus Information Technologies, Asheville, NC) using nose-point tracking. An object interaction was scored when the nose point was within 2 cm of the object center with the body axis of the mouse oriented towards the object. All preference scores were calculated from the test trial. “What” preference was calculated as the time exploring the two A objects subtracted from the exploration times of the two B objects (recent objects in corners 2 and 4), divided by the total times for all of the A and B objects. “When” memory preference was calculated as the exploration time for stationary A object (in corner 1) minus the exploration times of the two B objects (corners 2 and 4), divided by the total times with these three objects. “Where” preference score was calculated as the difference between the exploration time for the A object (which displaced object B in corner 3) minus the time with stationary A object (corner 1) divided by the total times with these two objects.

Twenty-four hours after completion of the W-W-W task, the mice were subjected to a seven-trial memory load test (Huffman et al., 2019; Miller-Rhodes et al., 2019). This test was conducted in the same arena as the W-W-W task, but without visual cues on the walls. Testing was performed over 7 consecutive test trials, with an approximate 60-90 sec inter-trial interval. To accommodate the increasing number of objects on each trial the first three trials were 3 min, trials 4-5 were 4 min, and trials 6 -7 were 5 min in length. On trial 1, one object was presented. With each subsequent trial a novel object (i.e., differing in configuration, color, and texture) was added in a random location to prevent the animal from predicting the new object based on the previous object’s placement. By the end of testing, all objects were equally spaced with no object being within 8-10 cm from the nearest object or wall of the test arena. Once the exploration time for a given trial, the mouse was placed into an empty mouse cage while the next object was placed in the arena before the mouse was reintroduced into the arena in the same location as before. Testing was filmed and scored using Noldus Ethovision 11 as in the W-W-W test. Exploration time with test objects was expressed as time spent in contact with the object/min to adjust for the varying lengths of the test-trials. Preference for the new object in each trial was expressed as the time spent with the new object minus the time spent with the remaining objects, divided by the sum of these times. The preference ratio ranged from −1 to +1, where positive scores indicated a preference for the new object, negative scores a preference for the previously introduced objects, and a score approaching “0” indicating no preference.

### Statistical Analysis

The behavioral data were analyzed with SPSS 25 (IBM SPSS, Chicago, IL), while all other data were analyzed with Prism 8 (GraphPad Software). The preference scores and time interacting with objects were analyzed with multivariate ANOVA (MANOVA). In the W-W-W test, the dependent variables were for the 3 different preference scores that were assessed simultaneously at testing. In the memory load test, the six different preference scores for trials 2-7 were the dependent variables. In all analyses the treatment condition was the fixed factor. *Post-hoc* tests were with Bonferroni corrected pair-wise comparisons. All non-behavioral data were analyzed by t-tests or ANOVA followed by Tukey’s multiple comparisons *post-hoc* tests. These data are presented as mean ± SEM. In all cases, a *P*<0.05 was considered statistically significant.

## Results

### Postoperative resolution of CNS inflammation and NLRP3 inflammasome activation

Failed resolution of CNS inflammation has been proposed as a driver of this condition in PNDs (Subramaniyan and Terrando, 2019), yet the role of ANXA1 in postoperative inflammation remains unknown. After orthopedic surgery, hippocampal expression of the endogenous pro-resolving mediator ANXA1 was significantly reduced, peaking at 24 hours (0.63 ± 0.13, *P* < 0.01) and almost fully resolving by 72 hours (1.13 ± 0.22, *P* < 0.01; **Fig. 1A**). This profile of impaired CNS inflammation resolution was paralleled by the expression of NLRP3 protein starting at 3 hours (0.81 ± 0.10, *P* < 0.01), peaking at 6 hours (1.75 ± 0.19, *P* < 0.001), remaining elevated at 24 hours (1.41 ± 0.22, *P* < 0.001), and returning to baseline by 72 hours (0.33 ± 0.22. *P* = 0.10), compared to naïve control (0.30 ± 0.11; *P* < 0.0001; **Fig. 1B)**.

**Figure 1.**
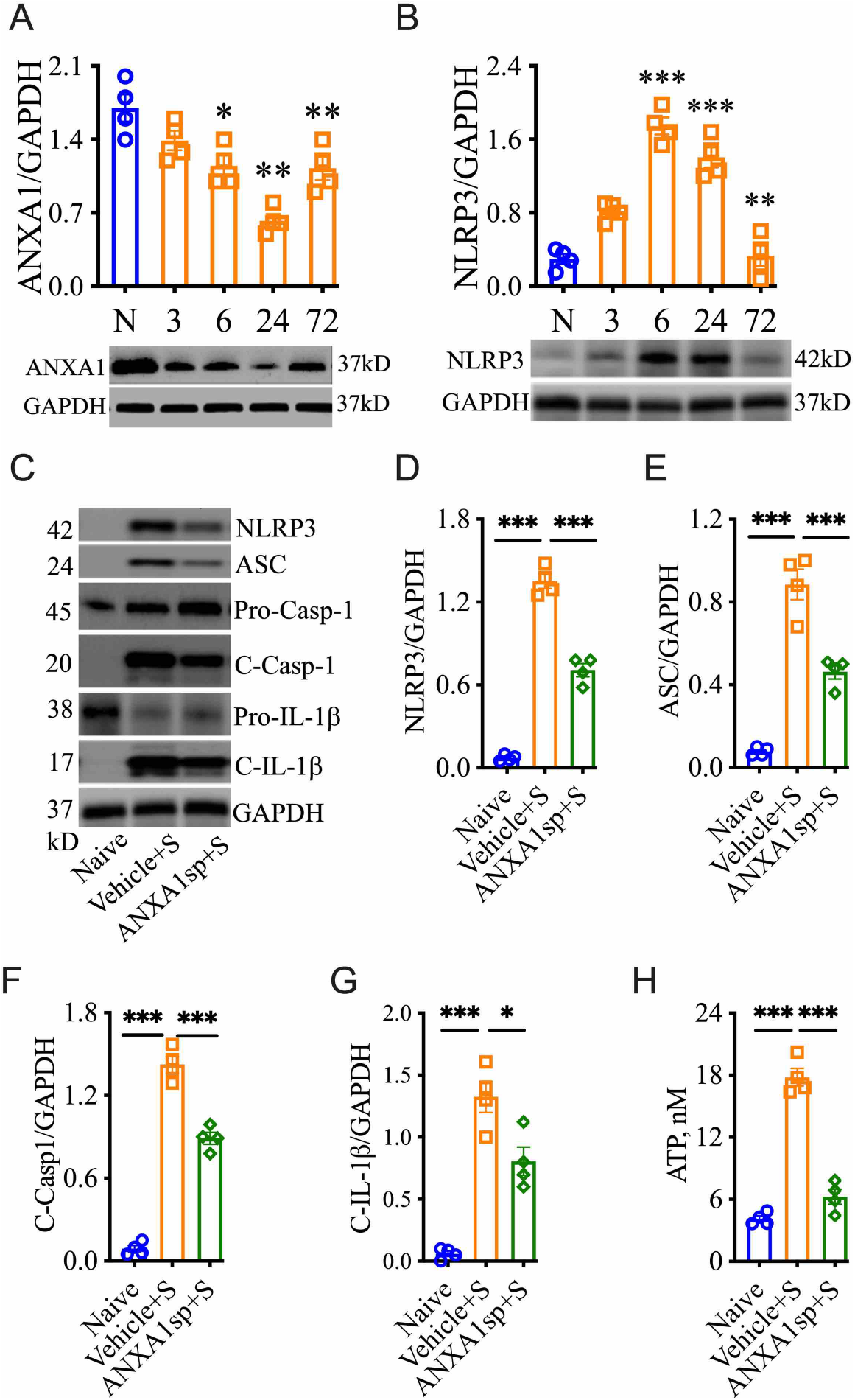
Activation of the NLRP3 inflammasome and resolution of inflammation in adult mice after orthopedic surgery. In adult mice subjected to orthopedic surgery, hippocampal expression of the endogenous pro-resolving mediator ANXA1 was significantly reduced in a time-dependent manner **(A)**. This reduction was accompanied by the induction of NLRP3 expression in the hippocampus (**B**). In adult mice treated with ANXA1sp (1mg/kg, i.p.) 30 minutes before surgery, postoperative NLRP3 inflammasome activation was significantly attenuated, as evidenced by Western blot analysis, particularly by attenuated levels of cleaved caspase-1 (**C-G**). ATP levels in hippocampal homogenate were also reduced, based on ELISA (**H**). Data are presented as mean ± SEM. *n* = 4. \**P* < 0.05, \*\**P* < 0.01, \*\*\**P* < 0.001 compared to naïve or vehicle controls. Analyzed by one-way ANOVA with Tukey’s multiple comparisons test.

Mice were treated with 1 mg/kg ANXA1sp (i.p.) or vehicle (0.5% DMSO in saline, i.p.) 30 minutes before surgery. After orthopedic surgery, the NLRP3 inflammasome was found to be activated in the CNS of vehicle-treated animals when compared to naïve controls (1.35 ± 0.10 vs 0.07 ± 0.02, *P* < 0.001; **Fig. 1C, D**), including increased ASC expression (0.89 ± 0.15 vs 0.08 ± 0.02, *P* < 0.001; **Fig. 1C, E**), activation of caspase-1 (1.43 ± 0.12 vs 0.09 ± 0.05, *P* < 0.001; **Fig. 1C, F**), and cleavage of IL-1β (1.33 ± 0.25 vs 0.06 ± 0.04, *P* < 0.001; **Fig. 1C, G**). Notably, NLRP3 inflammasome activation was effectively reduced in ANXA1sp-treated mice compared to vehicle-treated mice after surgery (0.71 ± 0.09 vs 1.35 ± 0.10, *P* < 0.001; **Fig. 1C, D**). Hippocampal expression of ASC (0.461 ± 0.07 vs 0.89 ± 0.15, *P* < 0.001; **Fig. 1C, E**), caspase-1 (0.89 ± 0.09 vs 1.43 ± 0.12, *P* < 0.001; **Fig. 1C, F**), and IL-1β (0.81 ± 0.23 vs 1.33 ± 0.25, *P* < 0.05; **Fig. 1C, G**) were all significantly inhibited in ANXA1sp-treated mice. Hippocampal levels of ATP were also upregulated after surgery in vehicle-treated animals compared to naïve controls (17.81 ± 1.72 vs 4.14 ± 0.58, *P* < 0.001; **Fig. 1H**). However, ATP was not induced in ANXA1sp-treated mice compared to naïve animals, while it was significantly reduced relative to vehicle-treated mice (6.26 ± 1.44 vs 17.81 ± 1.72, *P* < 0.001; **Fig. 1H**).

Morphologic changes in microglia have been reported in several models of surgery-induced cognitive impairments, including orthopedic surgery (Eckenhoff et al., 2020). Indeed, in our model of orthopedic surgery, we observed postoperative microglial morphologic changes in the hippocampus, with more stout (13.00 ± 4.69 vs 1.00 ± 1.00, *P* < 0.001; **Fig. 2A. B)**and fewer thin Iba-1-postive cells at 24 hours (4.80 ± 1.10 vs 21.60 ± 5.18, *P* < 0.001; **Fig. 2A, B**) indicating a pathological state compared to naïve controls. Tracing of these individual microglia further showed this stout morphology by decreased branch length with increased soma size after orthopedic surgery compared to thinner cells in naïve controls. Notably, postoperative microgliosis was improved in adult mice pretreated with ANXA1sp, with Iba-1 immunoreactivity shifted toward a more resting and ramified phenotype.

**Figure 2.**
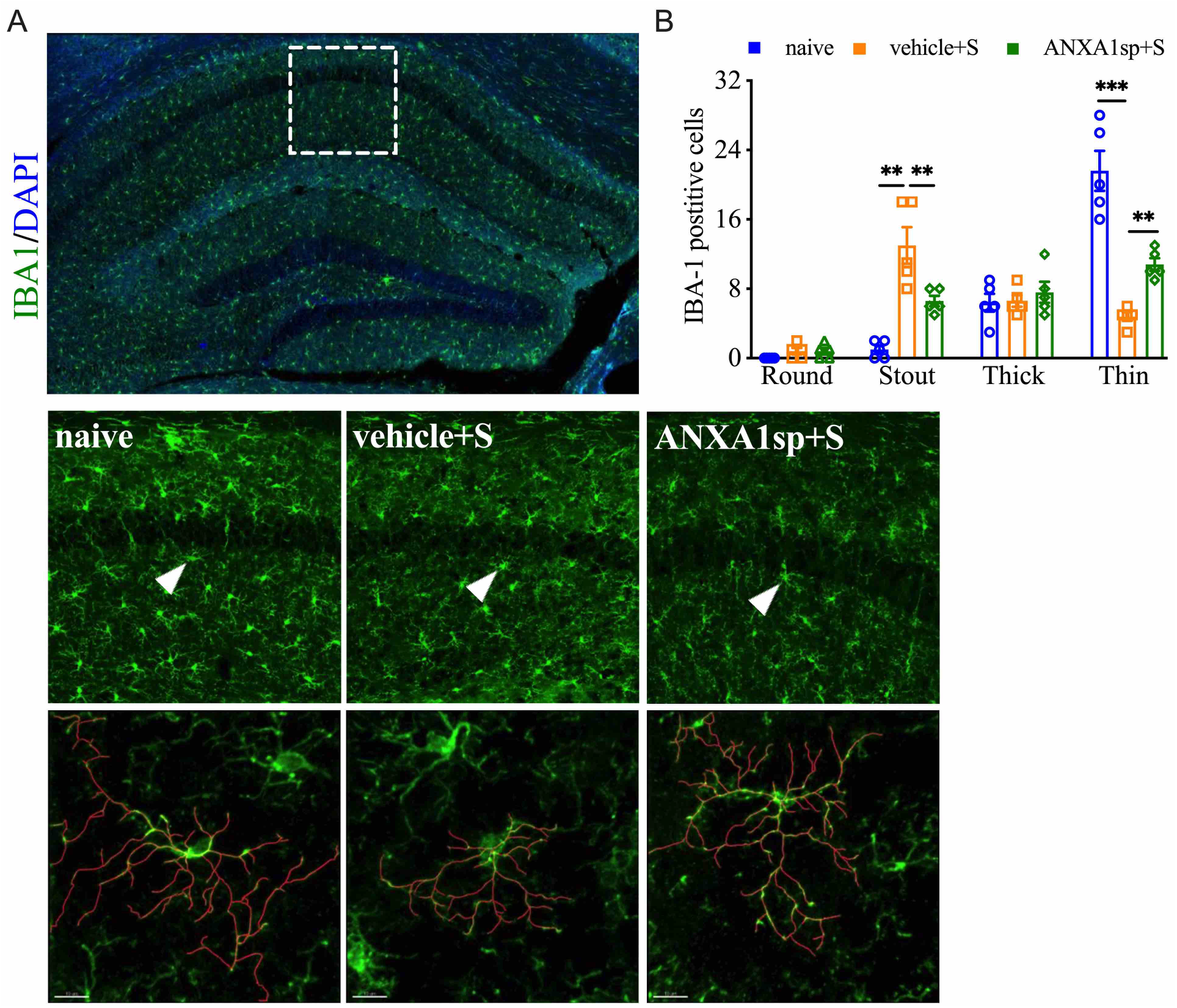
Effects of ANXA1sp on microglial morphology after surgery. Microglial morphologic changes were significantly curtailed 24 hours after orthopedic surgery in adult mice pretreated with ANXA1sp (**A, B**). Microglial morphology was quantified based on 4 morphologic subtypes: Round, Stout, Thick, and Thin; 3 representative areas/mouse were used for quantification and one single microglia was represented in the reconstruction (red indicates filament tracing). Data are presented as mean ± SEM. *n* = 5. \*\**P* < 0.01, \*\*\**P* < 0.001, compared to naïve or vehicle controls. Analyzed by one-way ANOVA with Tukey’s multiple comparisons test. Scale bar: 20 μm.

### Surgery-induced neuroinflammation and the NLPR3 inflammasome activation

To confirm the role of the NLRP3 inflammasome in surgery-induced neuroinflammation, *Nlrp3*^−/−^ and *Asc*^−/−^ adult mouse lines were used. *Nlrp3*^−/−^ mice remained protected from surgery-induced neuroinflammation. In fact, compared to wild-type mice, surgery in *Nlrp3*^−/−^ mice failed to upregulate expression of ASC (0.47 ± 0.57 vs 0.69 ± 0.57, *P* = 0.51), caspase-1 (0.26 ± 0.04 vs 0.28 ± 0.02, *P* = 0.44), and cleavage of pro-IL-1β (0.48 ± 0.04 vs 0.50 ± 0.06, *P* = 0.56), as shown by Western blot analysis (**Fig. 3A-D)**. Further, *Nlrp3*^−/−^ displayed no signs of reactive microgliosis at 24 hours after surgery, retaining thin and ramified morphologies (Round: 0.20 ± 0.45 vs 0.20 ± 0.45, *P* = 0.50; Stout: 0.6 ± 1.34 vs 0.20 ± 0.45, *P* = 0.29; Thick: 3.80 ± 1.79 vs 7.80 ± 5.81, *P* = 0.15; Thin: 21.20 ± 3.56 vs 19.40 ± 6.11, *P* = 0.34; **Fig. 3E, F**).

**Figure 3.**
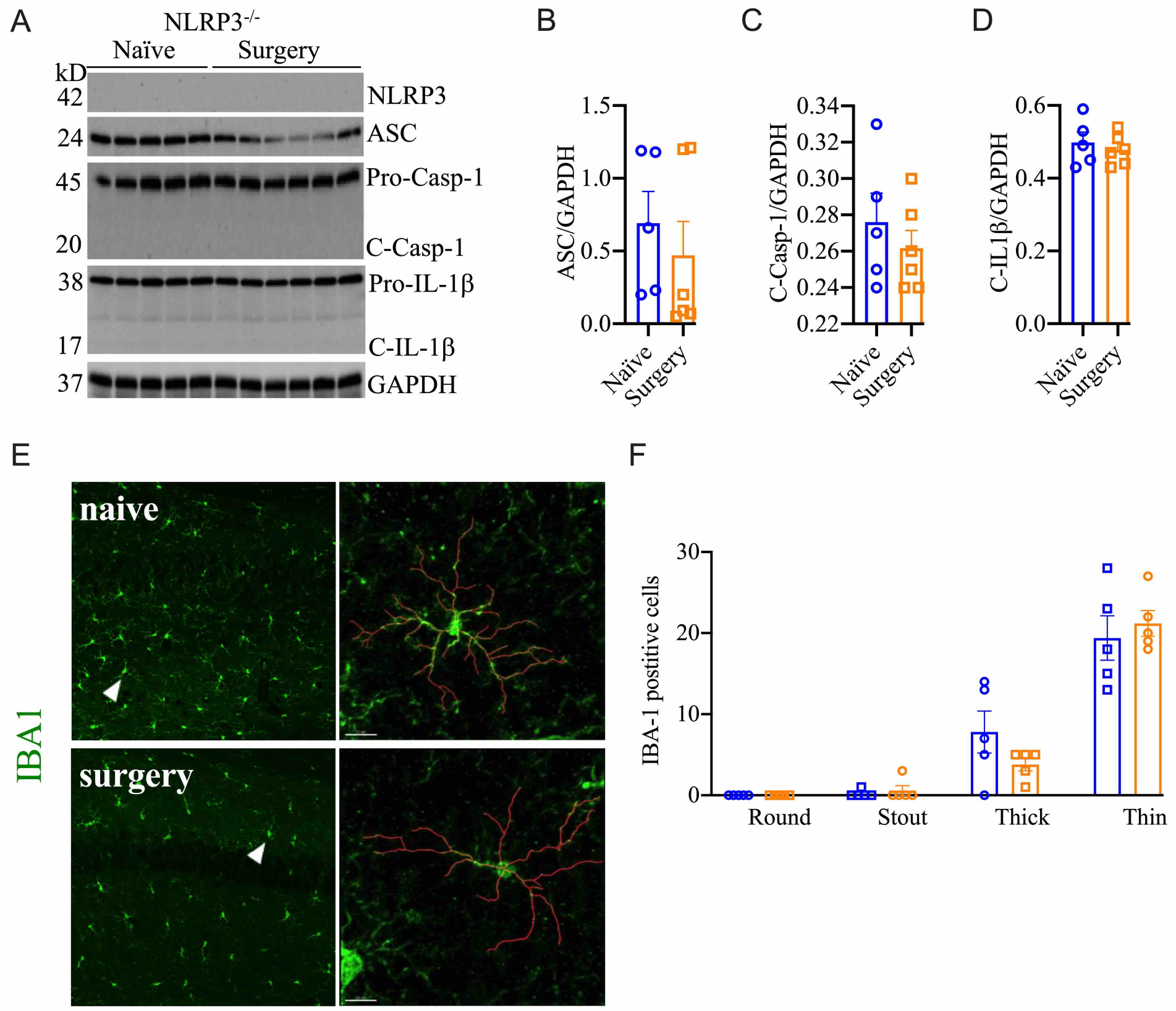
Postoperative neuroinflammation in *Nlrp3*^−/−^ mice. The NLRP3 inflammasome was not activated in the hippocampus of these animals at 24 hours post orthopedic surgery (**A-D**). Moreover, no changes in microglial morphology were observed in *Nlrp3*^−/−^ mice after orthopedic surgery, as evidenced by IBA-1 immunostaining (**E, F**). 3 representative areas/mouse were used for quantification and one single microglia was represented in the reconstruction (red indicates filament tracing. Data are presented as mean ± SEM. *n* = 5-6. Analyzed by Student *t* test and one-way ANOVA with Tukey’s multiple comparisons test. Scale bar: 20 μm.

Next, we evaluated *Asc*^−/−^ mice and found protection from surgical trauma that was similar to our findings in *Nlrp3*^−/−^ mice, with minimal effects on the NLRP3 complex, including NLRP3 (1.39 ± 0.05 vs 1.22 ± 0.14, *P* = 0.56), pro-caspase-1 (0.47 ± 0.05 vs 0.45 ± 0.03, *P* = 0.52), and cleavage of pro-IL-1β (0.85 ± 0.01 vs 0.80 ± 0.02, *P* = 0.13; **Fig. 4A-D**). To further evaluate the role of NLRP3 inflammasome activation in surgery-induced inflammation, we used ASC-citrine (ASC-cit) reporter mice. After surgery, ASC-cit mice displayed a significant increase in ASC speck formation, indicating the activated NLRP3 inflammasome complex, at 24 hours (42.80 ± 12.21 vs 21.60 ± 5.37, *P* < 0.01**; Fig. 4E, F**). These mice also showed signs of hippocampal microgliosis similar to C57BL6/J mice, with greater Iba-1 immunoreactivity and more stout cell bodies 24 hours after surgery (42.80 ± 12.21 vs 21.60 ± 5.37, *P* < 0.01; **Fig. 4E, F**). Notably, some of these ASC specks were co-localized with Iba-1-postive microglia (15.00 ± 1.00 vs 9.60 ± 1.40, *P* < 0.001; **Fig. 4E, G**).

**Figure 4.**
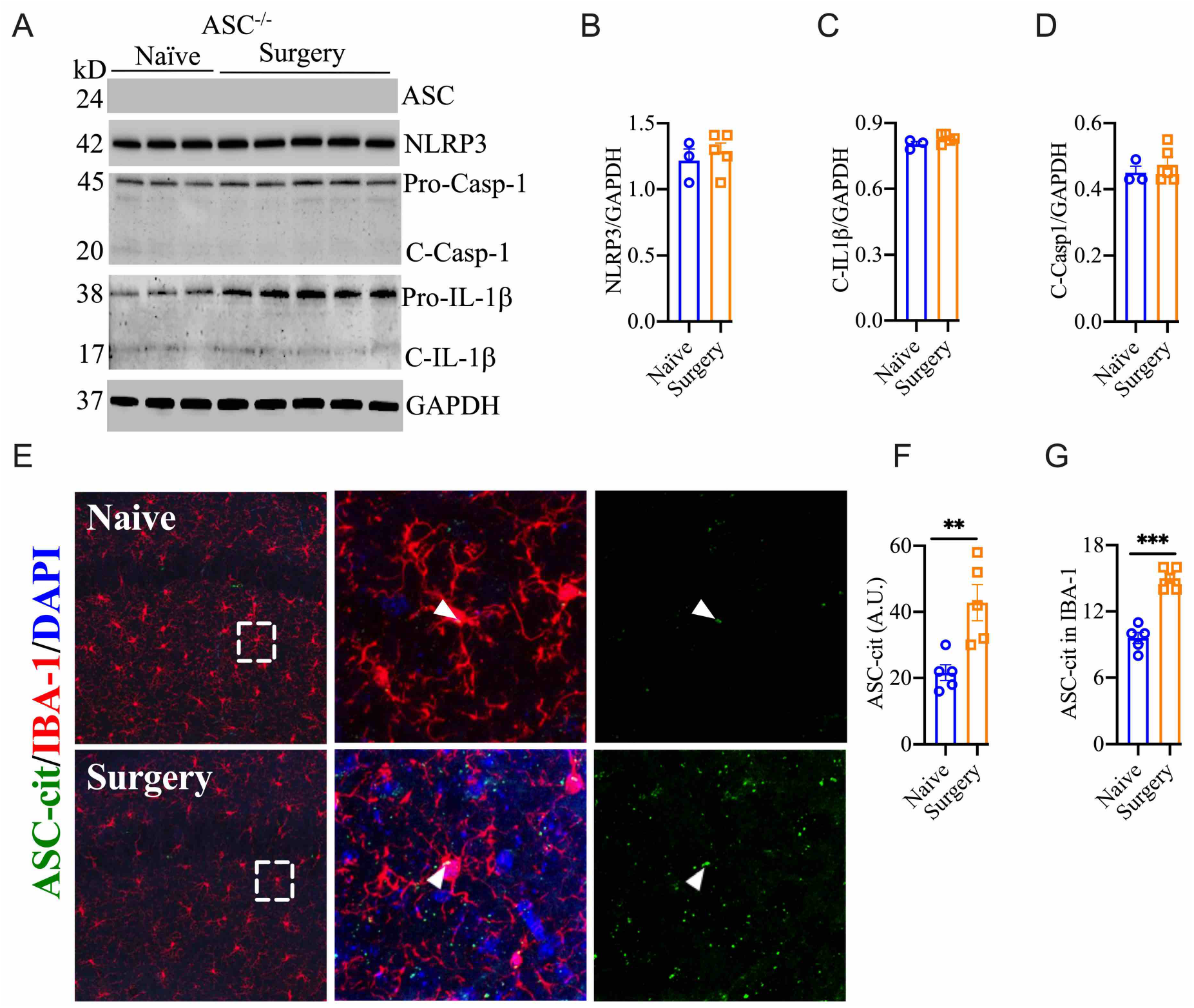
ASC speck formation after surgery. *Asc*^−/−^ showed no evidence of NLRP3 inflammasome activation in hippocampal lysate (**A-D**). ASC-cit reporter mice showed significant speck formation in the hippocampus, including co-expression with IBA-1-positive cells (**E-G**). 3 representative areas/mouse were used for quantification and one single microglia was represented in the reconstruction (red indicates filament tracing). Data are presented as mean ± SEM. *n* = 3-5. Analyzed by Student *t* test. Scale bar: 20 μm.

### PND-like behavior in adult mice pretreated with ANXA1sp

To study the functional effects of ANXA1sp, we evaluated cognitive responses that are affected in our PND-like model (Subramaniyan and Terrando, 2019). In the “What-Where-When” task (DeVito and Eichenbaum, 2010). mice were given free access to two sets of four objects (**Fig. 5A**). At testing, “What” memory was examined to determine whether the animal remembered the replacement of an object with another; “Where” memory analyzed if the animals recalled where an object had been previously located; and “When” memory reflected whether the mouse remembers the objects that were last investigated. For “What” memory, responses among the three groups (naïve, surgery plus vehicle injection, and surgery plus ANXA1sp treatment) were not statistically distinguished (**Fig. 5C**; **Table 1**). Nevertheless, marked differences in object preferences for the “Where” (**Fig. 5D**) and “When” (**Fig. 5E**) memory tests were evident. For “Where” memory the vehicle-treated mice with surgery showed a significant reduction in their abilities to identify the displaced object compared to the naïve group (*P*<0.011) and mice given ANXA1sp prior to surgery (*P*<0.038); these two latter groups had similar preferences for the displaced object. The “When” comparison was more difficult for the animals. Both the naïve and vehicle + surgery mice had similar moderate preference scores. By contrast, mice treated with ANXA1sp prior to surgery had significantly higher preference scores that the other two groups (*P*-values<0.052) animals. The higher preference scores for the ANXA1sp + surgery cannot be attributed to longer times spent with the objects since they had some of the lowest object interaction times across training and testing than any other group (**Fig. 5G**; **Table 2**). Indeed, no significant group differences in objet interaction times were discerned at training with the set A objects. At training with set B objects, mice treated with vehicle prior to surgery spent more time interacting with the objects compared to the naïve mice and those given ANXA1sp prior to surgery (*P*-values<0.051). Interestingly, at testing the vehicle + surgery and ANXA1sp + surgery mice showed spent less time exploring objects compared to the naïve animals (*P*-values<0.001).

**Table 1.** MANOVA results for “What”, “Where” and “When” preference scores.

| Test<sup>a</sup> | Degrees of Freedom | F-score | P Value |
| --- | --- | --- | --- |
| What | 2,27 | 2.762 | 0.081 |
| When | 2,27 | 5.845 | 0.008 |
| Where | 2,27 | 5.560 | 0.010 |

<sup>a</sup>N=10 mice/group (naïve, vehicle + surgery, ANXA1sp + surgery).

**Table 2.** MANOVA results for “What”, “Where” and “When” time spent exploring objects^a^.

| Test<sup>b</sup> | Degrees of Freedom | F-score | P Value |
| --- | --- | --- | --- |
| Set A Objects | 2,27 | 1.877 | 0.173 |
| Set B Objects | 2,27 | 5.391 | 0.011 |
| Test | 2,27 | 18.732 | 0.001 |

<sup>a</sup>Analysis is across the two training sessions with sat A and set B objects and at testing.
<sup>b</sup>N=10 mice/group (naïve, vehicle + surgery, ANXA1sp + surgery).

**Figure 5.**
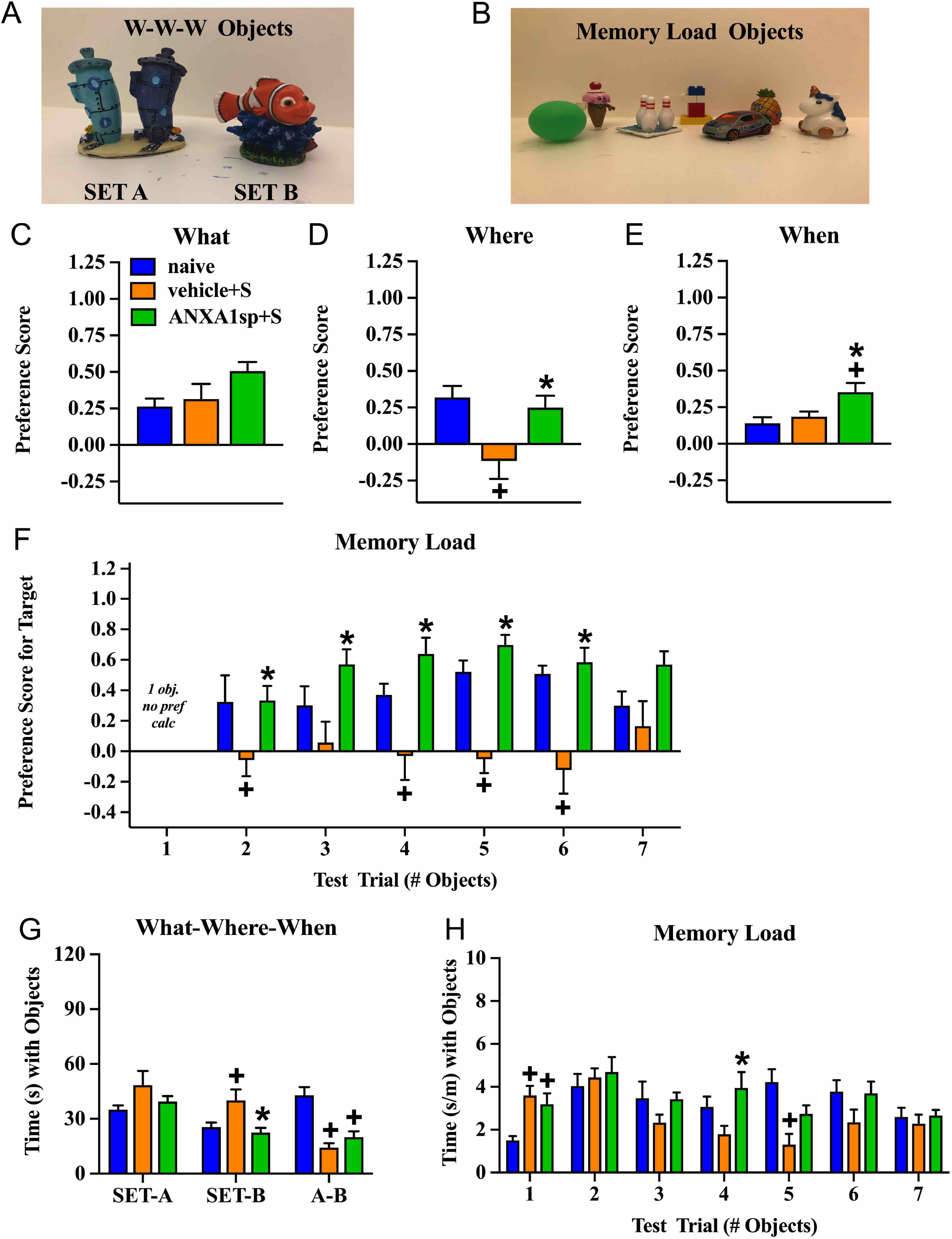
PND-like cognitive responses in adult mice pretreated with the vehicle or ANXA1sp. Example of Set A and Set B objects used in the “What-Where-When” memory task (**A**). Example of the seven unique objects used in the Memory Load task (**B**). Preference scores for “What” object recognition memory in the “What-Where-When” task (**C**). Preference scores for “Where” recognition memory for a displaced object in the “What-Where-When” task (**D**). Preference scores for “When” recognition memory for recent objects to previously investigated objects in the “What-Where-When” task (**E**). Preference scores for the novel object in the Memory Load test (**F**). The time (s) spent exploring Set A and Set B objects, and at testing in the “What-Where-When” task (**G**). Mean time (adjusted by trial length) that animals explored objects in the seven consecutive trials in the Memory Load test (**H**). N = 10 mice/treatment group for all tests. Results are shown as mean ± SEM and analyzed by MANOVA. \**P* < 0.05, compared to vehicle + surgery mice; +*P* < 0.05, compared to naïve animals.

In the memory load test naïve mice and those treated with ANXA1sp prior to surgery demonstrated an ability to discriminate the novel object even as memory load increased with up to six objects, while the mice treated with the vehicle prior to surgery were impaired (**Fig. 5F**; **Table 3**). Here, the vehicle + surgery mice showed significantly reduced preferences for the novel object compared to naïve group when 2 (*P*<0.052), 4 (*P*<0.051), 5 (*P*<0.001), and 6 (*P*<0.001) objects were present. Although performance by the mice in the naïve and ANXA1sp + surgery groups were similar, the treated group performed even better than the vehicle + surgery mice when 2 (*P*<0.057), 3 (*P*<0.018), 4 (*P*<0.001), 5 (*P*<0.001), and 6 (*P*<0.001) objects were presented. When time spent investigating objects across trails was examined, few group differences were observed (**Fig. 5H**; **Table 4**). However, on the first trial both treatment groups with surgery engaged in higher object exploration times than the naïve control (*P*-values<0.022). When 4 objects were present mice in the ANXA1sp + surgery group spent more time in object exploration than naïve mice (*P*<0.032). With 5 objects the naïve animals interacted with objects for a longer time than the vehicle + surgery mice (*P*<0.001). Together these results demonstrate that vehicle-treated tibia-fracture mice are markedly impaired in cognitive tasks compared to naïve mic, and that treatment with ANXA1sp prior to surgery prevents these cognitive impairments.

**Table 3.**
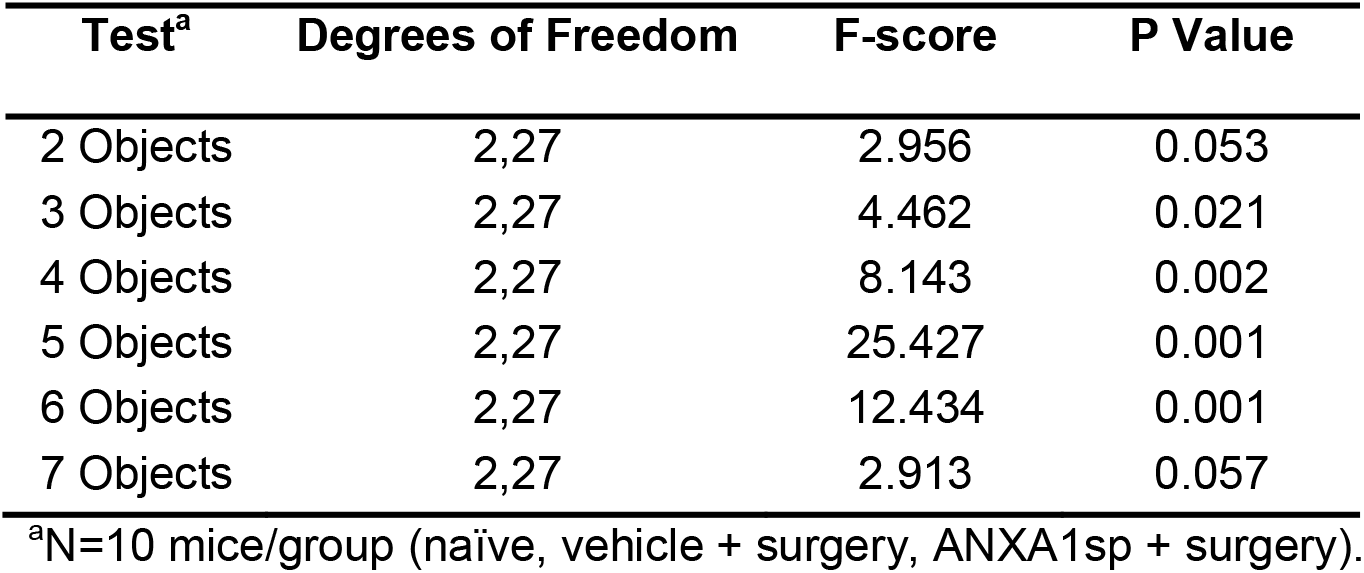
MANOVA results for memory load preference scores.

| Test<sup>a</sup> | Degrees of Freedom | F-score | P Value |
| --- | --- | --- | --- |
| 2 Objects | 2,27 | 2.956 | 0.053 |
| 3 Objects | 2,27 | 4.462 | 0.021 |
| 4 Objects | 2,27 | 8.143 | 0.002 |
| 5 Objects | 2,27 | 25.427 | 0.001 |
| 6 Objects | 2,27 | 12.434 | 0.001 |
| 7 Objects | 2,27 | 2.913 | 0.057 |

<sup>a</sup>N=10 mice/group (naïve, vehicle + surgery, ANXA1sp + surgery).

**Table 4.**
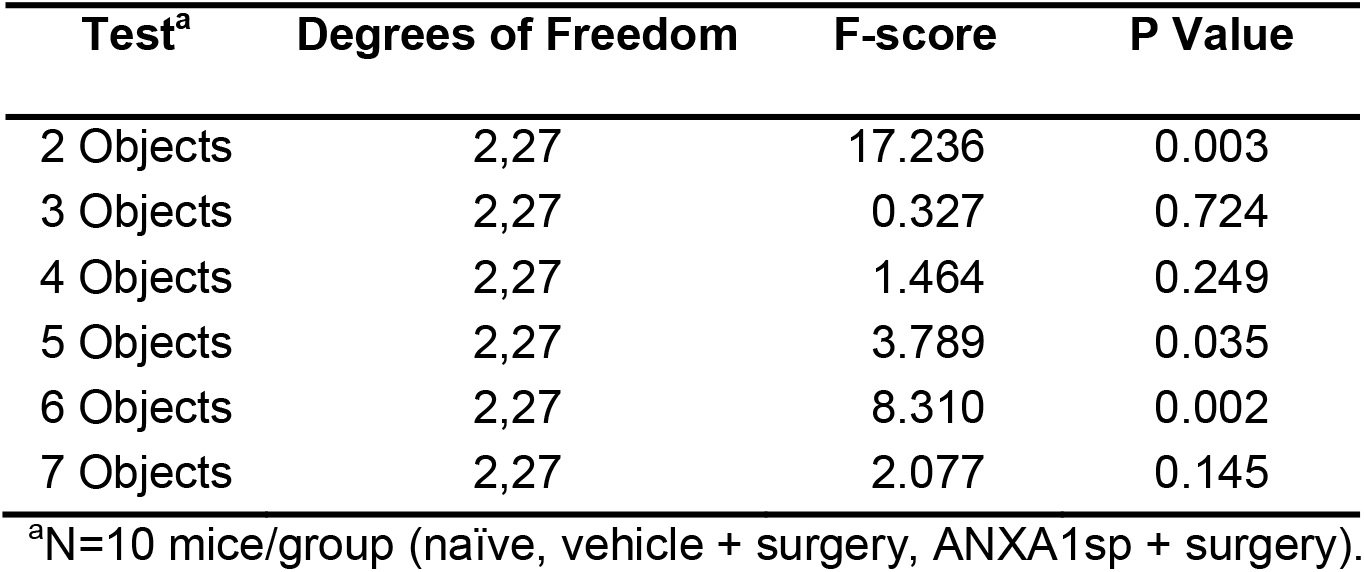
MANOVA results for memory load time spent exploring objects.

### Postoperative neuroinflammation in aged mice treated with ANXA1sp

Aging is a critical risk factor for many neurologic conditions, including PND (Lucin and Wyss-Coray, 2009). Thus, we administered the same therapeutic regimen of ANXA1sp in 18-22 month-old C57BL6/J mice, and assessed NLRP3 inflammasome activation and microgliosis. Compared to vehicle-treated adult mice, vehicle-treated aged mice showed a heightened, but similar, curve for NLRP3 activation after surgery, which peaked at 6 hours (2.48 ± 0.14, *P* < 0.0001), remained elevated at 24 hours (2.00 ± 0.30, *P* < 0.0001), and resolved to baseline by 72 hours (0.98 ± 0.11, *P* = 0.28) compared to naïve controls (0.70 ± 0.19); *P* < 0.0001; **Fig. 6A**). Notably, more NLRP3 activation was found in aged mice compared to adult mice after surgery [R^2^ = 0.98; F(1,3) = 130.5]; *P* = 0.0014; **Fig. 6B**]. Western-blot analysis showed increased levels of NLRP3 in the hippocampus (1.95 ± 0.16 vs 0.32 ± 0.11, *P* < 0.001; **Fig. 6C-G**). This exacerbated NLRP3 inflammasome activation was significantly blunted in aged mice pretreated with ANXA1sp (1.39 ± 0.08 vs 1.95 ± 0.16, *P* < 0.001; **Fig. 6C, D**), affecting the expression of ASC (ANXA1sp vs vehicle: 0.85 ± 0.23 vs 1.79 ± 0.34, *P* < 0.001; **Fig. 6C, E**), activation of caspase-1 (ANXA1sp vs vehicle: 1.51 ± 0.07 vs 2.18 ± 0.20, *P* < 0.001; **Fig. 6C, F**), and levels of IL-1β (ANXA1sp vs vehicle: 1.77 ± 0.15 vs 2.50 ± 0.36, *P* < 0.001; **Fig. 6C, G)**. Further, hippocampal ATP was retained at baseline levels in aged mice pretreated with ANXA1sp vs vehicle (17.56 ± 2.51 vs 29.6 ± 4.51, *P* < 0.001; **Fig. 6H**).

**Figure 6.**
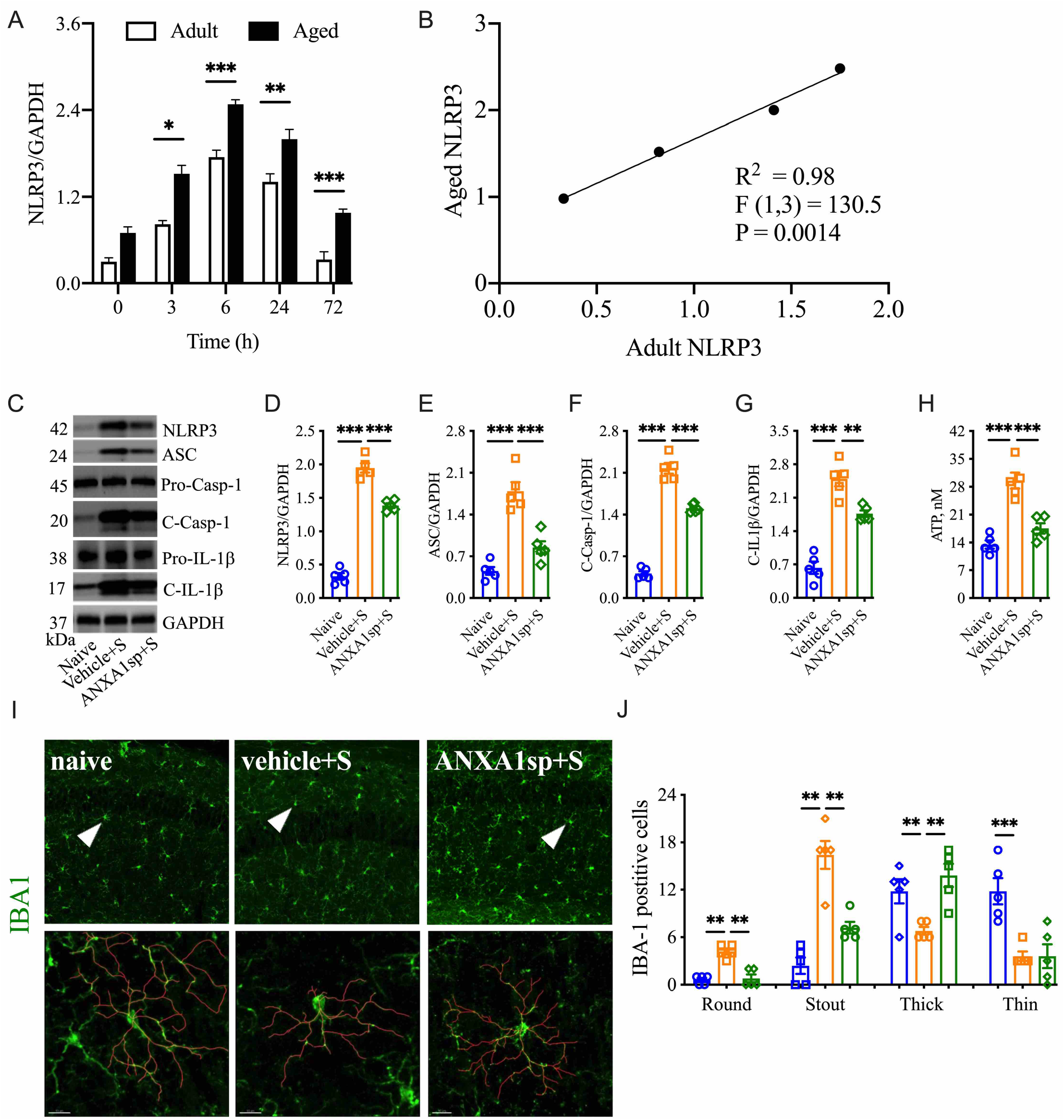
Postoperative neuroinflammation in aged mice. The hippocampal NLRP3 inflammasome complex was significantly activated in a time-dependent manner in aged mice after orthopedic surgery **(A-B)**. NLRP3 inflammasome activation (**C-G**) and hippocampal ATP expression (**H**) were attenuated in aged mice treated with ANXA1sp (1 mg/kg, ip) 30 minutes before orthopedic surgery (same dose and timepoint as adult mice). Microglial activation was also improved 24 hours after surgery (**I, J**). 3 representative areas/mouse were used for quantification and one single microglia was represented in the reconstruction (red indicates filament tracing). Data are presented as mean ± SEM. *n* = 5. \*\**P* < 0.01, \*\*\**P* < 0.001, compared to naïve or vehicle controls. Analyzed by one-way ANOVA with Tukey’s multiple comparisons test. Scale bar: 20 μm.

We next evaluated microglial activation, and observed a similar age-dependent priming in Iba-1-postive cells (*P* < 0.0001; **Fig. 6I, J**). Surgery triggered more stout (16.40 ± 3.97 vs 2.40 ± 2.30, *P* < 0.01) and round (4.20 ± 0.84 vs 0.60 ± 0.55, *P* < 0.01) microglia in the hippocampus with reduced branch length compared to naïve controls. The round microglia were hardly visible in adult mice at the same time point (*P* < 0.01). Importantly, aged mice pretreated with ANXA1sp showed significantly improved microglial pathology, especially soma enlargement and swelling, 2 key features of CNS dyshomeostasis (Streit et al., 2004). However, arborizations were not restored in the aged brain of pretreated mice (*P* < 0.001).

## Discussion

Here, we studied the relationship between resolution agonists, particularly ANXA1 signaling, and NLRP3 inflammasome activation in mediating postoperative neuroinflammation and PND-like behavior. We found that preoperative treatment with a small peptide derived from the N-terminal region of human ANXA1 (ANXA1sp) attenuates NLRP3 inflammasome activation in adult mice after orthopedic surgery, and thereby reduces microgliosis and behavioral deficits. Notably, these anti-neuroinflammatory effects were also observed in aged mice, representing a potential next step toward translating resolution agonists into perioperative clinical trials.

Dysregulated immunity has become a key pathologic hallmark of virtually every neurologic disorder including PNDs. Surgical trauma triggers a systemic inflammatory response that can impair microglial function, and thus contribute to sickness behavior and memory decline. This postoperative cytokine storm has been well documented across several rodent models as well as in surgery patients. In our earlier work, we discovered a role for IL-1β signaling in mediating surgery-induced neuroinflammation. Mice treated with IL-1 receptor antagonist (IL-1RA) or lacking IL-1R showed no signs of neuroinflammation or microglial activation (Cibelli et al., 2010). Subsequent studies in rats after abdominal surgery further defined the importance of *central* IL-1β signaling in contributing to PND-like behavior. In fact, only intracisternal administration of IL-1RA was effective in reducing behavioral impairments and neuroinflammation after surgical trauma (Barrientos et al., 2012). Studies in older adults with delirium after hip fracture repair confirmed the presence of high levels of IL-1β in cerebrospinal fluid (CSF), suggesting that IL-1 may represent a valuable biomarker for identifying and possibly treating patients at risk for PNDs (Cape et al., 2014). Indeed, IL-1β signaling has been described as a key mediator of sickness behavior (fever, HPA axis activation, and depression)(Anforth et al., 1998) as well as microglial priming during aging, and overall neuroinflammation (reviewed in (Konsman et al., 2002)). However, despite seminal contributions in this field, little is known about the regulation of mature IL-1β in PNDs.

IL-1β maturation is critically regulated by the NOD-like receptor protein 3 (NLRP3) inflammasome. Canonical activation of the NLRP3 inflammasome involves recruitment of adapter proteins, including apoptosis-associated speck-like protein containing a caspase recruitment domain (ASC), and caspase-1, which orchestrate the maturation and secretion of IL-1β and IL-18 (Schroder and Tschopp, 2010). Canonically, generation of mature IL-1β requires 2 signals, the production of pro-IL-1β by activating transcription factors, such as NF-κB, and maturation of IL-1β and its release by pyroptosis induced by NLRP3 inflammasome activation, stimulated by factors such as ATP (Bauernfeind et al., 2009). Surgery triggers NF-κB activation *via* multiple pathways including danger-associated molecular patterns (DAMPs), released during the acute-phase response following aseptic surgical trauma.

DAMPs, such as HMGB1, are potent activators of NF-κB and the subsequent cytokine storm, which has been well described in this model of orthopedic surgery (Terrando et al., 2010; Vacas et al., 2014; Yang et al., 2019). In fact, disabling NF-κB activation in this model prevents microglial activation, further highlighting the critical role of this gene in regulating the innate immune response to surgery (Terrando et al., 2011). Absence of key immune receptors, such as MyD88 and TLR4, prevents cytokine release and surgery-induced microglial activation (Terrando et al., 2010; Lu et al., 2015). In the current study, we found that NLRP3 and ASC knock-out mice remained protected from postoperative neuroinflammation, indicating an upstream mechanism for NF-κB activation. Treatment with ANXA1sp downregulated the expression of key inflammasome components, including ASC and NLRP3, and levels of IL-1β after surgery in both adult and aged mice (**Fig. 7**).

**Figure 7:**
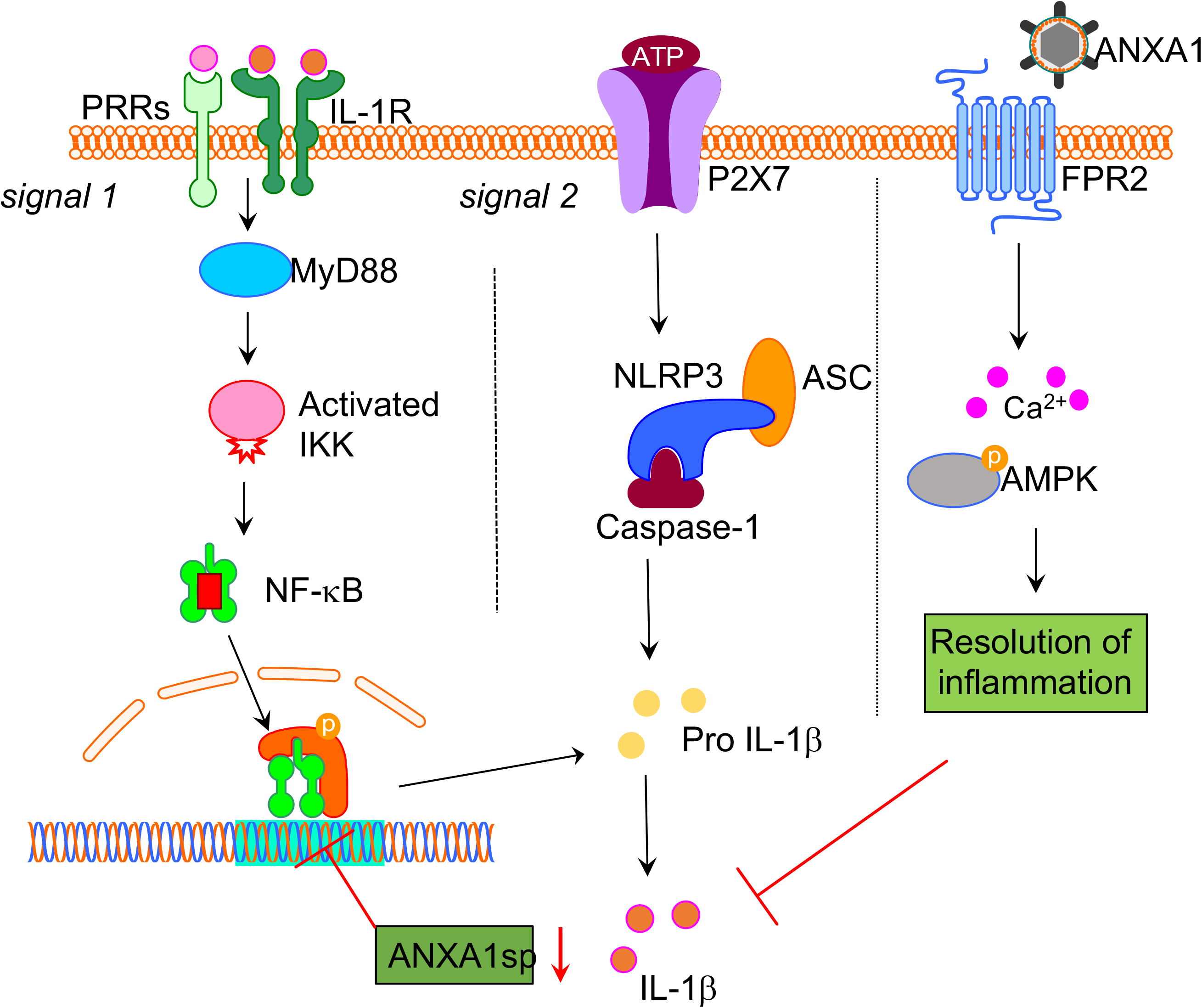
Proposed NLPR3 signaling mechanism in postoperative neuroinflammation. Aseptic surgical trauma induces a cytokine storm that engages several pattern recognition receptors (PRRs), including the interlukin-1 (IL-1) receptor. Downstream signaling from these receptors relies on adaptor proteins such as MyD88, which eventually activates NF-κB to perpetuate the inflammatory response. Parallel to this process, surgery also elevated levels of ATP in the hippocampus. In combination with NF-κB activation this induced the NLRP3 inflammasome and ASC specks formation in microglia, thus contributing to postoperative neuroinflammation. Surgery also impaired the expression of hippocampal Annexin-A1 (ANXA1) that is critically involved in inflammation-resolution and can be regulated through our biomimetic peptide. Within this framework we found that treatment with ANXA1sp regulated NLRP3 expression and levels of IL-1β postoperatively. Given the established elevation of IL-1β in preclinical and clinical biofluids of subjects with postoperative neurocognitive disorders harnessing this pathway may provide safer strategies to treat ensuing neuroinflammation.

Notably, we found evidence of NLRP3 inflammasome activation in microglia after surgery with formation of “ASC specks”, which result from recruitment and polymerization of ASC into ~1-μm structures (Stutz et al., 2013). Specks are not unique to activation of the NLRP3 inflammasome, but also occur with activation of NLRP1b, NLRC4, and AIM2 (Chan and Schroder, 2020). The elevated ATP that we measured in whole hippocampal homogenate after surgery may be responsible for the microglial response, thus leading to speck formation. Indeed, extracellular ATP is a well-known activator of microglial processes after injury (Davalos et al., 2005) and an activator of the NLRP3 inflammasome, and surgery may hijack their neuroprotective functions leading to a neurotoxic environment and PNDs. Additional work using more detailed approaches is needed to further characterize the microglial response and the signaling events following peripheral surgery.

Previous work described a key role for microglial activation in PNDs. PET imaging provides evidence of microgliosis in postsurgical patients (Forsberg et al., 2017). Here, we found that preoperative administration of ANXA1sp reduced morphologic changes associated with microglial activation in the hippocampus. These effects may result from direct modulation of resident microglia since ANXA1 is expressed on these cells, and the small peptide can easily cross the blood-brain barrier (BBB) (Sheikh and Solito, 2018). Although we did not evaluate BBB opening in the present study, it is important to note that orthopedic surgery impairs BBB integrity and the expression of tight junctions such as claudin-5 and aquaporin-1, thus facilitating infiltration of peripherally injected compounds into the CNS (Yang et al., 2019; Wang et al., 2020). Indeed, this breach also allows systemic factors such as IgG and fibrinogen as well as inflammatory myeloid cells such as macrophages to enter the brain parenchyma and contribute to neuroinflammation (Terrando et al., 2011; Velagapudi et al., 2019). Preventing infiltration of CCR2-positive monocytes after surgery reduces microgliosis and PND-like behavior (Degos et al., 2013). Given that ANXA1 is a key regulator of endothelial function and BBB integrity, it is plausible that these neuroprotective effects are mediated by limiting leukocyte trafficking at the neurovascular unit (Cristante et al., 2013). In fact, ANXA1 peptidomimetics such as Ac2-26 exert potent effects on the vasculature by limiting neutrophil adhesion and emigration, but without impairing cell rolling (Lim et al., 1998). ANXA1 is also highly expressed on macrophages, facilitating efferocytosis of apoptotic neutrophils and overall resolution of inflammation (Scannell et al., 2007; Dalli et al., 2012). The pro-resolving effects of ANXA1sp may target this critical step at the level of the BBB, although our peptide also exerts potent anti-inflammatory effects on systemic cytokines (Zhang et al., 2017). Thus, ANXA1sp effects on the CNS may result from an overall dampening of the pro-inflammatory milieu rather than, or in addition to, a direct effect on the BBB.

ANXA1 is a 37-kDA protein that is a member of a larger annexin superfamily of calcium-dependent phospholipid-binding proteins that signal *via* G protein-coupled receptors and formyl peptide receptor 2 (FPR2) (Walther et al., 2000), thus inhibiting phospholipase A_2_ (PLA_2_) and downstream production of bioactive lipid mediators derived from arachidonic acid release (Flower and Blackwell, 1979). Notably, the tripeptide sequence of ANXA1sp (Ac-Gln-Ala-Trp) at the N-terminus represents a critical domain for signaling *via* FRP2 and exert anti-inflammatory effects (Movitz et al., 2010). ANXA1 also facilitates resolution of inflammation by acting on the lipoxin A4 receptor (ALXR) to downregulate polymorphonuclear neutrophil recruitment at the site of inflammation (Perretti et al., 2002). These actions are important because they limit the synthesis of eicosanoids such as prostaglandins, thromboxanes, prostacyclins, and leukotrienes, which are upregulated in the CNS of patients after peripheral surgery (Buvanendran et al., 2006). ANXA1 signaling *via* ALXR was recently shown to support regeneration of skeletal muscle after injury (McArthur et al., 2020), which may translate to the surgical setting. These regulatory effects of ANXA1 and many other pro-resolving mediators provide unique opportunities to apply resolution pharmacology to the clinical setting, given their known safety profile.

In summary, we have found a previously unrecognized action of an ANXA1 peptidomimetic on the NLRP3 inflammasome complex, which resulted in reduced microglial activity and PND-like behavior after surgery. Although further work is needed to determine whether ANXA1sp directly binds to the NLRP3 inflammasome complex, our findings provide a novel target for regulating postoperative neuroinflammation and cognitive disorders during aging.

## Acknowledgments

This work was in part supported by the National Institutes of Health (NIH) R01AG057525, a School of Medicine voucher for studies conducted in the Mouse Behavioral and Neuroendocrine Analysis Core Facility, and Duke Anesthesiology Departmental funds. Some of the behavioral experiments were conducted with equipment and software purchased with a North Carolina Biotechnology Center grant. We would like to thank Mr. Christopher Means for his support in behavioral testing and Kathy Gage, BS (Duke University Medical Center) for editorial assistance.

## Notes

**Conflict of interest statement:** ZZ and QM are coinventors on patents filed through Duke University on the therapeutic use of ANXA1sp. Other authors declare no competing financial interests.

### Competing Interest Statement

ZZ and QM are coinventors on patents filed through Duke University on the therapeutic use of ANXA1sp. Other authors declare no competing financial interests.

